# Acupuncture triggers earlier recovery from ischemic stroke than sham needling in a rat model

**DOI:** 10.1101/2024.07.14.601555

**Authors:** Wuxian He, Hongtu Tang, Jia Li, Xiaoyan Shen, Chenrui Li, Huafeng Liu, Weichuan Yu

## Abstract

Acupuncture, a traditional Chinese medical treatment that has been practiced for over 2,000 years, is widely used around the world [1]. However, its efficacy and distinction from random stimulation are still being questioned [2, 3]. Over the years, many studies have reported either favorable, neutral or even skeptical outcomes regarding the treatment effect of acupuncture on diverse ailments [4–7]. The major question behind this controversy is whether acupuncture is different from random needle insertion and whether its efficacy can be attributed to the placebo effect [8, 9]. Here, we use micro-positron emission tomography (microPET) imaging in a randomized controlled animal study to show that acupuncture facilitates faster recuperation in comparison to sham acupuncture and blank control. Based on the microPET imaging of subjects undergoing daily acupuncture over two weeks’ duration, we dynamically monitored the metabolic activity levels in different brain regions and found that both acupoint and non-acupoint stimulation could improve ischemic stroke recovery. This finding is consistent with previous reports that both acupuncture and sham needling show a positive effect in the treatment of diseases [4, 5, 8]. More importantly, we further found that rats receiving acupuncture at Baihui (GV20) and Shuigou (GV26), two commonly used acupoints for stroke rehabilitation based on the concept of the meridian system, showed earlier recovery effects than rats receiving sham needling treatment. This difference mainly appeared in regions involved in motor control and was validated by a balance beam walking test. Our findings, in conjunction with a recent electroacupuncture study that revealed a neuroanatomical pathway to mediate the vagal-adrenal anti-inflammatory axis [10], provide quantitative evidence supporting the specificity of acupoints in acupuncture therapy.

## Introduction

Acupuncture is one of the oldest treatment methods in traditional Chinese medicine, first recorded in the *Huangdi Neijing* (*The Yellow Emperor’s Internal Classic*) around 100 BC [11]. It has been employed by Chinese people for generations and in the West since the 20th century [12]. The core principle of acupuncture treatment is the hypothesis that the human body comprises acupoints that are interconnected via meridian channels [13], and needling at distinct acupoints can modulate different tissues and organs through the meridians, thereby alleviating pain or treating disease [14, 15]. Despite having demonstrated efficacy in treating diverse practical symptoms, acupuncture has encountered skepticism within Western medicine due to a lack of scientific evidence supporting its underlying mechanisms [2, 16].

Existing research outcomes on the effectiveness of acupuncture are disparate. Supporting evidence includes the finding that needle insertion can prompt the mobilization of endogenous opioid peptides from the central nervous system, resulting in acupuncture-induced analgesia [17]. Additionally, a recent study revealed a neuroanatomical basis underlying the anti-inflammatory effect driven by the vagal-adrenal axis triggered through electroacupuncture stimulation at the Zusanli acupoint [10, 18]. Nonetheless, investigations examining the efficacy of acupuncture in treating chronic low back pain [8] and chronic knee pain [5], preventing migraines [4], and rehabilitating stroke patients [19, 20] have shown that acupuncture is not superior to sham acupuncture or standard therapy. Furthermore, it has been shown that needling at non-acupoint sites elicits the same effect as needling at acupoints [21]. Given that the majority of these studies were clinical trials in which patients were fully aware of the needle insertion, and assessments were based on patient questionnaires or physician evaluations, acupuncture has been regarded as having a potent placebo effect [22].

In this study, we have devised a preclinical randomized controlled animal trial to investigate the therapeutic efficacy of acupuncture on rats with ischemic stroke (Fig. 1a). By employing animal subjects under anesthesia, we can mitigate the impact of the placebo effect. Moreover, we have included an adequate number of subjects to minimize experimental bias and enhance reproducibility. Each rat was subjected to brain ischemia induction, followed by daily acupuncture plus microPET imaging for ten days in a two-week duration. The functional neuroimaging data enables us to monitor metabolic activity in various brain regions, thereby studying the specific effects of acupoints in stroke rehabilitation. The findings exhibited a high degree of consistency with the behavioral assessment scores obtained via the balance beam walking test.

**Fig. 1.**
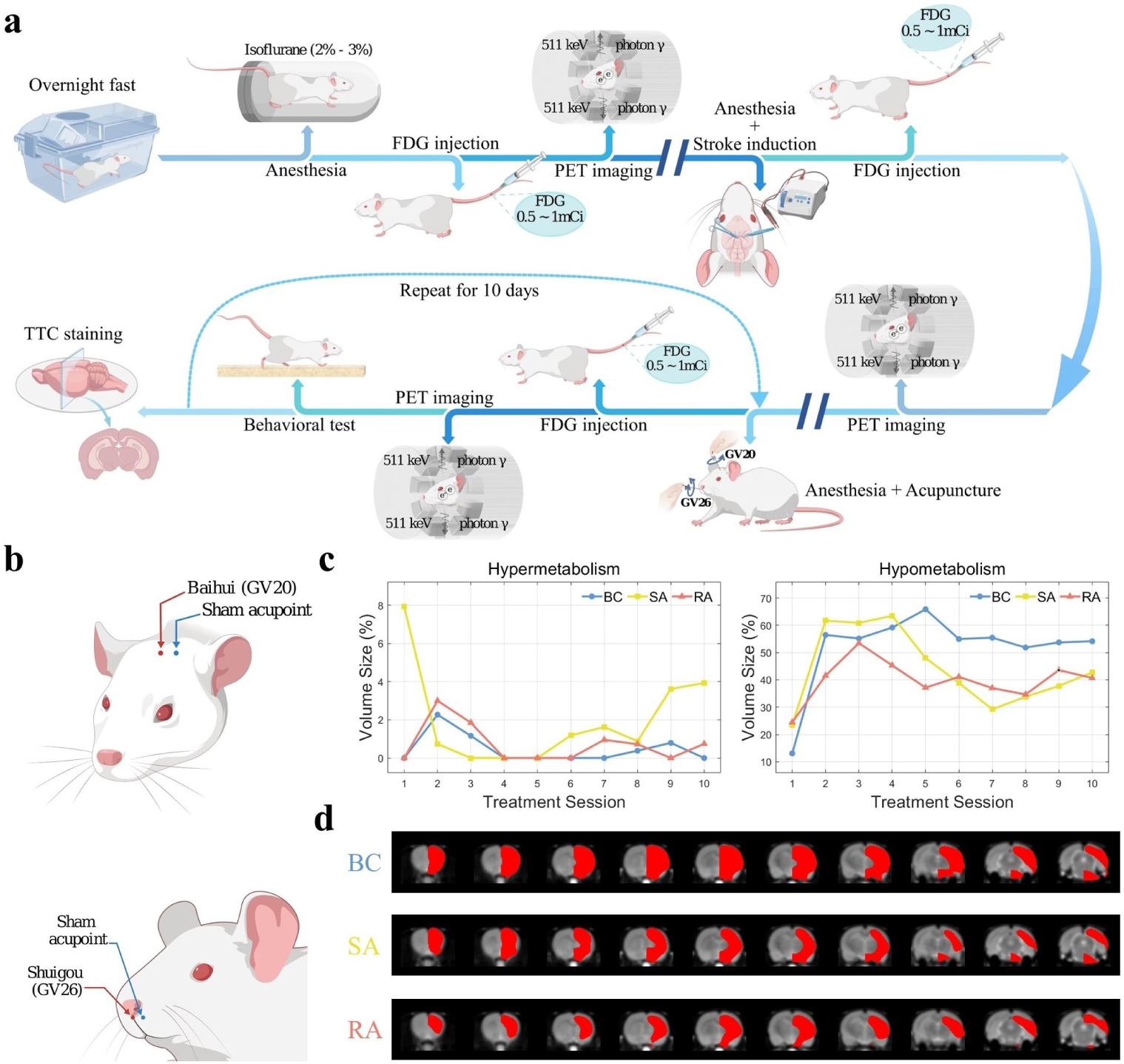
Graphic illustration of the experimental workflow and the initial findings. **a**, The flowchart of the animal experiment. Rats from the RA and SA groups received needling at real and sham acupoints, respectively, while rats from the BC group received no treatment. MicroPET imaging served as a primary tool for assessing brain functional activity, while behavioral tests and TTC staining were employed as validation methods. **b**, The locations of the real acupoints (GV20 and GV26) and sham acupoints. The two sham acupoints are positioned 5 mm to the right of GV20 and GV26, correspondingly. **c**, The percentages of hypermetabolic (left) and hypometabolic (right) volume sizes in the ipsilateral right brain across the entire right hemisphere. Voxel-wise analysis was performed to determine the significance by comparing each treatment scan with the MCAO scan. A one-tailed paired *t*-test was conducted for each voxel, with hypermetabolism and hypometabolism accounting for each direction of the one-tailed test. Significant voxels were identified with false discovery rate (FDR) corrected at a level of *q* = 0.01 using the Benjamini-Hochberg (BH) procedure [30]. The results of using a more conservative Bonferroni multiple correction show a similar trend (Extended Fig. 10). The volume size was obtained by counting the number of hypermetabolic/hypometabolic voxels over the size of the entire right brain. **d**, A visualized example of the hypometabolic locations after the fifth treatment session. The hypometabolic voxels (in red) are overlaid on the brain template (background), and ten coronal slices with uniform spacing are presented. The leftmost slice is situated at 4.00 mm to the anterior commissure, while the rightmost slice is located at 7.09 mm to the anterior commissure.

### Both needling at acupoints and non-acupoints exhibit efficacy for metabolic recovery

The study compares the therapeutic efficacy of real acupuncture (RA) applied at GV20 and GV26 with that of sham acupuncture (SA) and with a blank control (BC) group that received no treatment. GV20 and GV26 are two commonly targeted acupoints in stroke research [23], located at the vertex of the dorsal midline and 1 mm below the nasal tip, respectively [24]. The SA followed the same procedure as RA but at selected non-acupoints on the right side of GV20 and GV26, each at a 5 mm distance (Fig. 1b). The rats first underwent middle cerebral artery occlusion (MCAO) to induce brain ischemia and were randomly assigned to a treatment group thereafter. Each subject had MCAO surgery on a Monday, and animals from the RA and SA groups received their first treatment on the same day (4 h following MCAO), continuing until the next Friday, for ten consecutive working days (as depicted in Fig. 1a).

[18F]-fludeoxyglucose (FDG) microPET images were acquired before the MCAO, after the MCAO, and after each treatment session. Voxels exhibiting significant metabolic changes in the ipsilateral hemisphere were computed based on hypermetabolism (i.e., increased voxel intensity) and hypometabolism (i.e., decreased voxel intensity) by subtracting each treatment scan from the MCAO scan. The trends of the time series of hypermetabolism show a concave shape, with greater FDG uptake observed at the beginning and end of the study, and lower uptake in the middle (Fig. 1c left). This pattern of hyper-uptake is consistent with previous studies, which have reported that increased FDG uptake is usually observed during the acute phase after MCAO and after one week of brain ischemia [25]. Acute hypermetabolism typically occurs in the penumbra 24 h after the onset of brain ischemia, which coincides with the timing of the second treatment scan in our experiment. The cause of this acute hypermetabolism is likely the up-regulation of glucose transporters or elevated glycolysis in the viable penumbra tissue [26]. On the other hand, delayed hypermetabolism occurs 7 d after ischemia, which is primarily due to neuroinflammation [27].

Regarding the ischemic tissue fate, research has demonstrated that hypermetabolism exhibits a limited connection with cerebral dysfunction. In contrast, ischemic cores displaying reduced glucose uptake possess a strong correlation with tissue infarction and neurological deficits [28]. The findings of our study indicate that hypometabolism exhibited a rapid expansion from the second treatment session (day 2) onwards and persisted in more voxels throughout the entire treatment period compared to hypermetabolism (Fig. 1c right). Consequently, to investigate the sustained therapeutic impact of acupuncture on brain injury following cerebral ischemia, we concentrated on scrutinizing regions with metabolic decline. The temporal patterns of the hypometabolic volumes reveal that rats without any treatment (i.e., the BC group) exhibited the most substantial hypo-uptake volumes overall, whereas rats subjected to acupuncture at real acupoints and sham acupoints both showed significantly reduced volume sizes (Fig. 1c right and d). This suggests that needle stimulation, at either real or sham acupoints, exerts a therapeutic impact on stroke rehabilitation and yields a comparable effect to noninvasive brain stimulation techniques [29]. Interestingly, we further observe that the curve of the RA group shows an earlier decline than that of the SA group (Fig. 1c right), suggesting that rats receiving needling at real acupoints may have experienced faster recovery.

### The specificity of real acupoints in triggering earlier recovery

To investigate the specific effect of acupuncture on the recovery pattern in the RA group (Fig. 1c right), a more comprehensive delineation of regions of interest (ROIs) was undertaken. This was accomplished by using a volumetric rat brain atlas (i.e., the Waxholm Space atlas) consisting of 222 ROIs [31] and aligning it to our microPET images. Voxel-wise analysis was conducted on individual regions to determine the hypometabolic volume. The findings reveal that RA led to a faster shrinking of hypometabolism than SA in motor control areas such as the motor cortex and somatosensory cortex.

The area most closely related to motor control is the motor cortex. Signals of body movement were generated from the motor cortex, which is a part of the neocortex located in the frontal lobe (Fig. 2a and b). We derived the hypometabolic volume in the primary motor cortex (M1) and secondary motor cortex (M2) for each treatment day by comparing the corresponding PET images with the scans taken after MCAO. The dynamic trend thus obtained reveals the neuronal metabolic deficit of each area on a daily basis for all three groups. Starting from day 4 onwards, rats belonging to the RA group exhibited a significant decrease in hypometabolic volumes within M1 compared to those in the BC group (Fig. 2c), suggesting the start of metabolic recovery. In contrast, rats from the SA group did not display a significant reduction in hypometabolic volumes until day 6. Meanwhile, the hypometabolic volumes in the RA group dropped to zero and remained at that level until the last day, whereas the hypometabolic volumes in the SA group dropped to zero on day 7 and rebounded on the eighth day, eventually reaching the same level as the BC group. Comparable trends can be observed in M2 (Fig. 2d), where rats in the RA group demonstrated a significant reduction in hypometabolic volumes from day 4 onwards, with the magnitude remaining below 60% after day 6. On the other hand, the SA group exhibited a similar degree of hypometabolic volume to the BC group during the initial six days, followed by a sudden metabolic recovery on day 7. Subsequently, the SA group exhibited a rebound effect, reaching a higher level compared to the BC group by the end of the experiment. Localization analysis of the hypometabolic volume reveals that the recovery of metabolism in both M1 and M2 originated from the outer regions (Fig. 2e and f), which may be attributable to neurovascular remodeling in the penumbra area mediating the switch in the tissue from injury to repair [32].

**Fig. 2.**
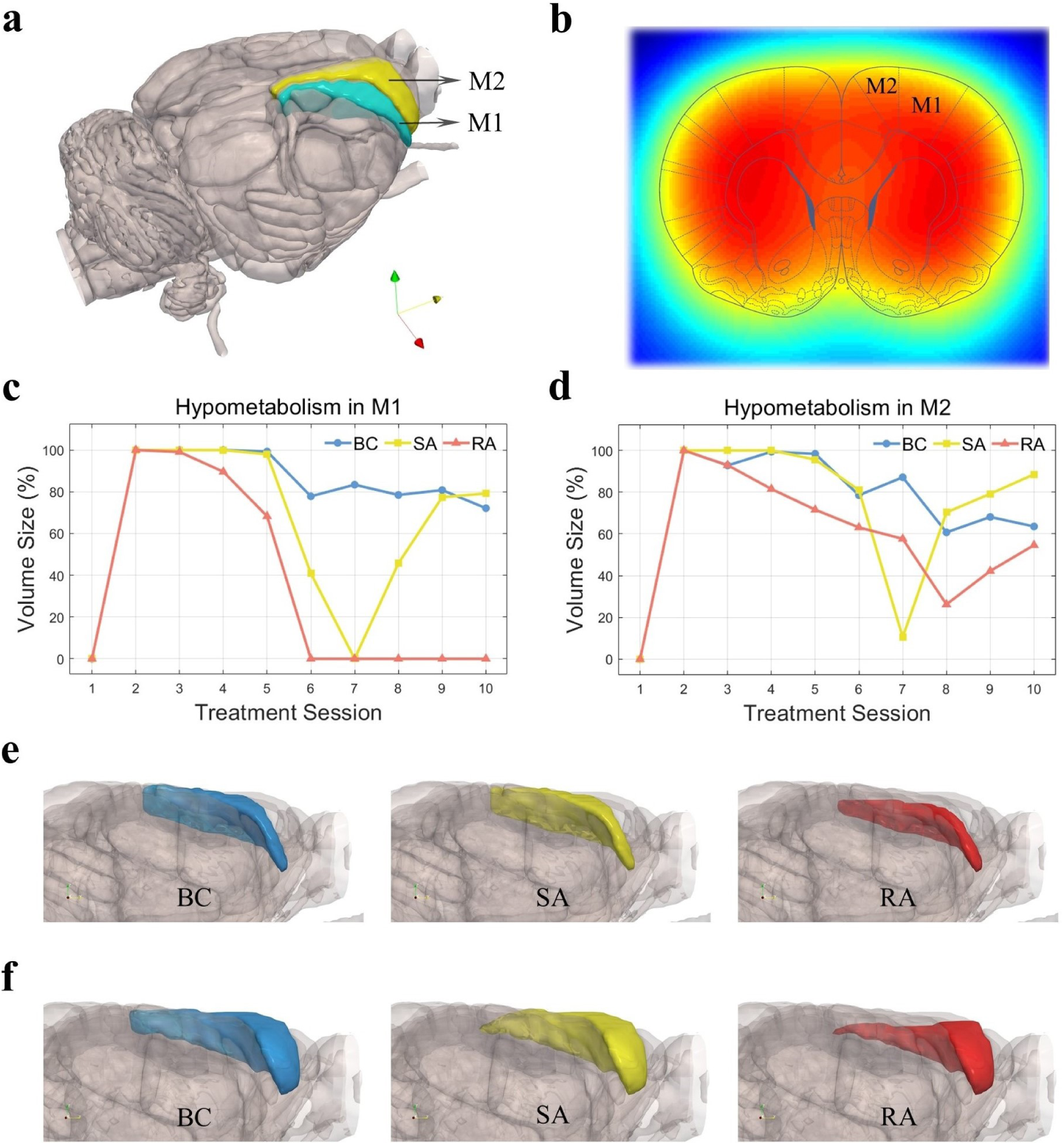
Hypometabolic volumes in the motor area. **a**, The locations of the primary motor cortex (M1) and secondary motor cortex (M2) in the 3D atlas. **b**, The locations of M1 and M2 in a 2D coronal slice overlaid on the FDG microPET images. **c**, **d**, The ratio of voxels exhibiting hypometabolism in the ipsilateral M1 (**c**) and M2 (**d**) over their original volume sizes, respectively. Voxel-wise analysis was performed to determine the significance by comparing each treatment scan with the MCAO scan. A one-tailed paired *t*-test was conducted for every voxel within each region, and significant voxels were identified with FDR corrected at a level of *q* = 0.01 using the BH procedure. Sample sizes in each treatment group (BC, SA, and RA) are *n* = 52, 57, 55 (day 1); *n* = 55, 59, 58 (day 2); *n* = 51, 55, 55 (day 3); *n* = 51, 54, 51 (day 4); *n* = 52, 51, 44 (day 5); *n* = 58, 51, 49 (day 6); *n* = 57, 53, 51 (day 7); *n* = 50, 55, 47 (day 8); *n* = 56, 56, 52 (day 9); and *n* = 53, 56, 50 (day 10). The fluctuation in sample sizes on different days was attributed to the exclusion of microPET images showing an incorrect field of view (FOV) or distortion. **e**, The locations of hypometabolic volumes in the ipsilateral M1 on day 5. Results shown from left to right represent the BC, SA, and RA groups, respectively. **f**, The locations of hypometabolic volumes in the ipsilateral M2 on day 5.

In addition to the motor cortex, another cortical region that is closely associated with a majority of body movements is the somatosensory cortex, particularly the primary somatosensory cortex (S1). The S1 is located within the neocortex and plays an important role in receiving sensory information and processing motor signals for movement control [33]. We investigated the metabolic levels within sub-regions of S1, and the results reveal that early recovery signals in the RA group mainly existed in the forelimb representation (S1FL), hindlimb representation (S1HL), and trunk representation (S1Tr). These three areas, identified through electrophysiological mapping [34], are interconnected and are adjacent to the M1 (Fig. 3a and b). The hypometabolic volume size of these three areas showed a similar trend, with a significant reduction in both the RA and SA groups in the mid-term of the treatment period compared to the BC group (Fig. 3c-e). Furthermore, similar to the early recovery signal observed in M1 and M2, the hypometabolic volume sizes in these three areas also demonstrated an earlier decrease in the RA group, after day 4, compared to the SA group. The early recovered cells were once again located in the periphery (Fig. 3f-h), comprising the penumbra of S1. Meanwhile, within the SA group, the rats showed a rebound of hypometabolism around days 7-8 after the volume dropped to zero, surpassing even the level observed in the BC group at the end of the experiment (Fig. 3c-e). In contrast, although the rats in the RA group also showed a rebound of hypometabolic volume in S1HL and S1Tr, the rebound trend was reversed on the last day and dropped back to zero. The overall pattern of hypometabolic volumes in S1FL, S1HL, and S1Tr was highly consistent with that observed in M1 and M2. This suggests a strong correlation in functional recovery among these areas and highlights the potential for neuroplasticity in facilitating rehabilitation after stroke [29].

**Fig. 3.**
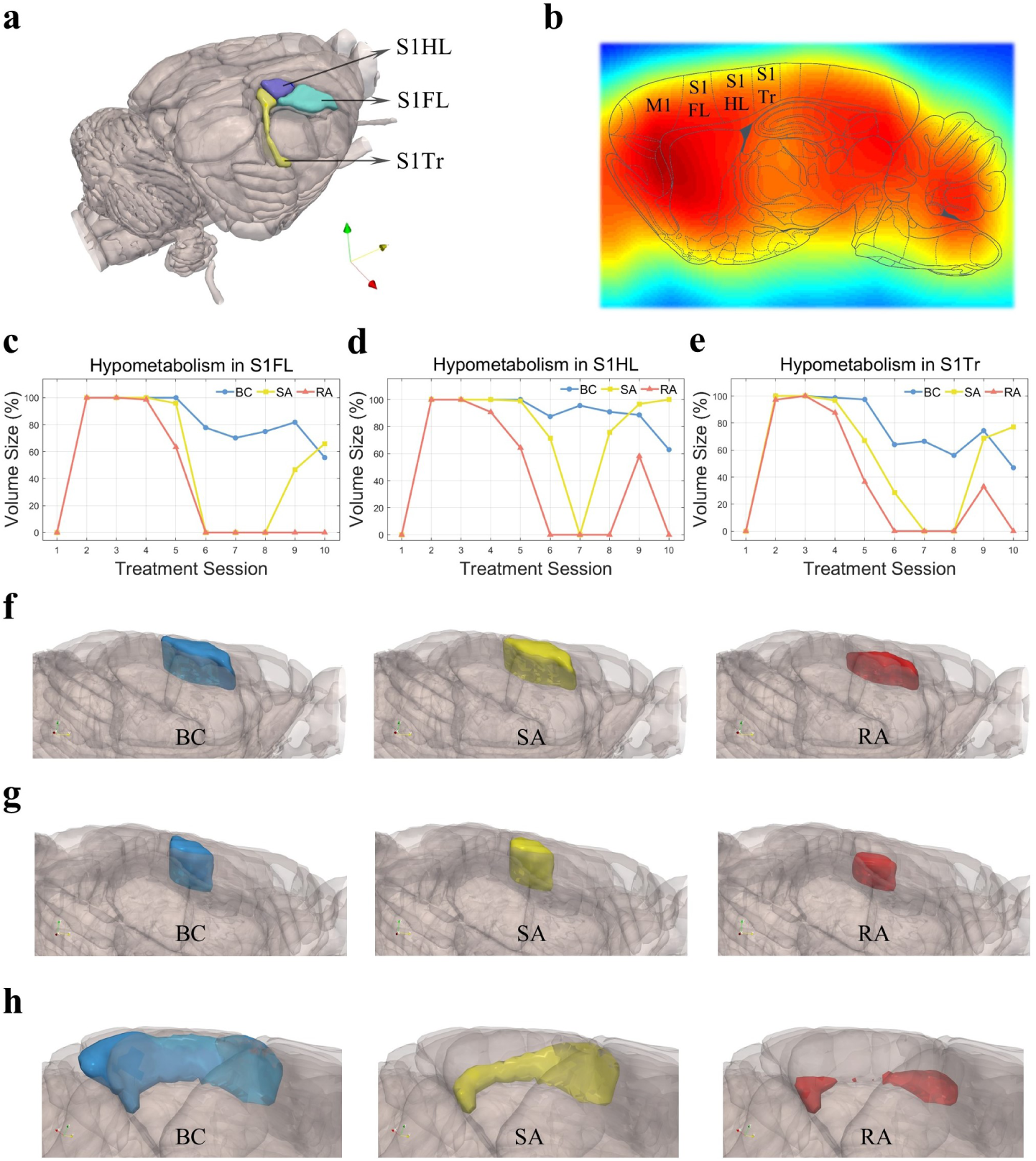
Hypometabolic volumes in the primary somatosensory area. **a**, The locations of the primary somatosensory cortex: forelimb representation (S1FL), hindlimb representation (S1HL), and trunk representation (S1Tr) in the 3D atlas. **b**, The locations of S1FL, S1HL and S1Tr in a 2D coronal slice overlaid on the FDG microPET images. **c** -**e**, The ratio of voxels exhibiting hypometabolism in the ipsilateral S1FL (**c**), S1HL (**d**) and S1Tr (**e**) over their original volume sizes, respectively. Calculations and sample sizes are the same as those described in Fig. 2c. **f**, The locations of hypometabolic volumes in the ipsilateral S1FL on day 5. Results shown from left to right represent the BC, SA, and RA groups, respectively. **g**, The locations of hypometabolic volumes in the ipsilateral S1HL on day 5. **h**, The locations of hypometabolic volumes in the ipsilateral S1Tr on day 5.

### Assessment of brain lesion and motor impairment

The brain suffers from reduced cerebral blood flow and oxygen supply after focal ischemia, which increases the risk of damage to the affected tissue. Acute-stage ischemic brain lesions were identified using T2-weighted MRI imaging, conducted 2 h after the MCAO procedure. The MRI scans were aligned to the Waxholm Space atlas to locate the lesion sites. (Fig. 4a). The results show that the MCAO procedure in this experiment resulted in tissue injuries in the bottom right region of the cerebral cortex (indicated by the red crosshairs in Fig. 4a) two hours after the ligation. According to the Waxholm Space atlas, this region within the brain lesion corresponds to parts of the piriform cortex, endopiriform nucleus, insular cortex, and claustrum. These areas were previously identified in prior studies as infarcted areas following MCAO [35, 36], and this was further confirmed by TTC staining conducted 24 h after MCAO in this experiment (Fig. 4b).

**Fig. 4.**
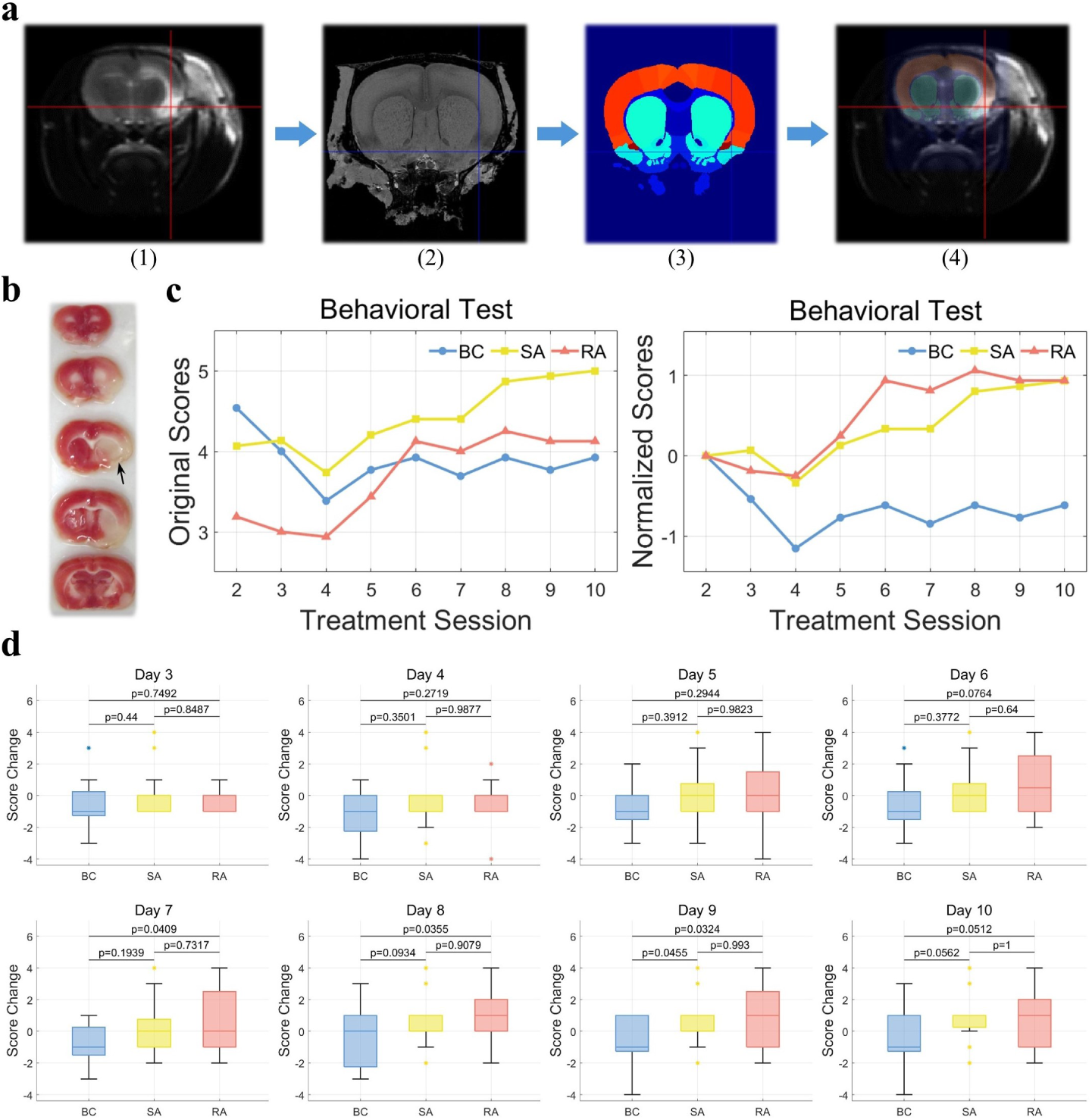
Validation of the ischemic model and motor recovery effect. **a**, Ischemia-induced brain lesion was indicated by the crosshairs in one example of a T2-weighted MRI scan (1). The acquired MRI scans were aligned to the MRI atlas (2) provided in [31], which had been previously registered to the Waxholm Space atlas (3). Precise localization of the brain lesion (4) could be achieved based on the volumetric atlas. **b**, A TTC staining example of rat brain sections performed 24 h post brain ischemia. The white area represents tissue infarction, which corresponds to the injured area shown in the MRI images in (a), as indicated by the black arrow. **c**, Left, original behavioral scores (averaged) from the balance beam walking test with smaller scores indicating more severe motor deficit. Right, baseline-adjusted behavioral scores (averaged) computed by subtracting the scores of each rat from their initial scores obtained during the first test. The first behavioral test was carried out on day 2 due to an overload of experimental procedures on the first day. **d**, Comparisons of the normalized behavioral scores from (c) on a daily basis. The box plot shows the minimum, 25th percentile, median, 75th percentile, and maximum values of the scores. Dotted data are outliers having values more than 1.5 times the interquartile range away from the bottom or top of the box. One-way ANOVA with a post-hoc Tukey’s test; *n* = 13, 15, 16 for the BC, SA, and RA groups, respectively.

A balance beam walking test was conducted daily from the second day of treatment onwards to assess the motor deficit of the rats in the BC/SA/RA groups. A seven-point grading scale was adopted to evaluate the rats’ ability to navigate the balance beam, with a score of 1 indicating the most severe motor deficit and a score of 7 indicating the least severity (see Methods and Extended Table 1). Since each rat obtained different scores in the first test (Fig. 4c left), the scores were adjusted by subtracting their initial test scores, establishing a baseline in order to compare the behavioral score trends throughout the treatment sessions (Fig. 4c right). The normalized scores represented changes relative to the initial results, with positive values indicating improvement in motor functioning and negative values indicating a worsening of motor ability. Despite the relatively wide range of normalized behavioral scores observed within each group, rats in the RA group demonstrated statistically significant superior performance compared to rats in the BC group from day 7 onwards (Fig. 4d). In contrast, rats in the SA group exhibit statistically better performance relative to the BC group only after day 9, despite no significant distinction between the RA and SA groups throughout the treatments. This earlier restoration of motor ability in the RA group than in the SA group by two days is consistent with the earlier decrease in hypometabolic volumes estimated using microPET images for rats in the RA group relative to rats in the SA group. A strong negative correlation also exists between the size of the hypo-uptake volume in the ipsilateral hemisphere and the motor ability assessed through the beam walking test (Extended Fig. 5).

**Fig. 5.**
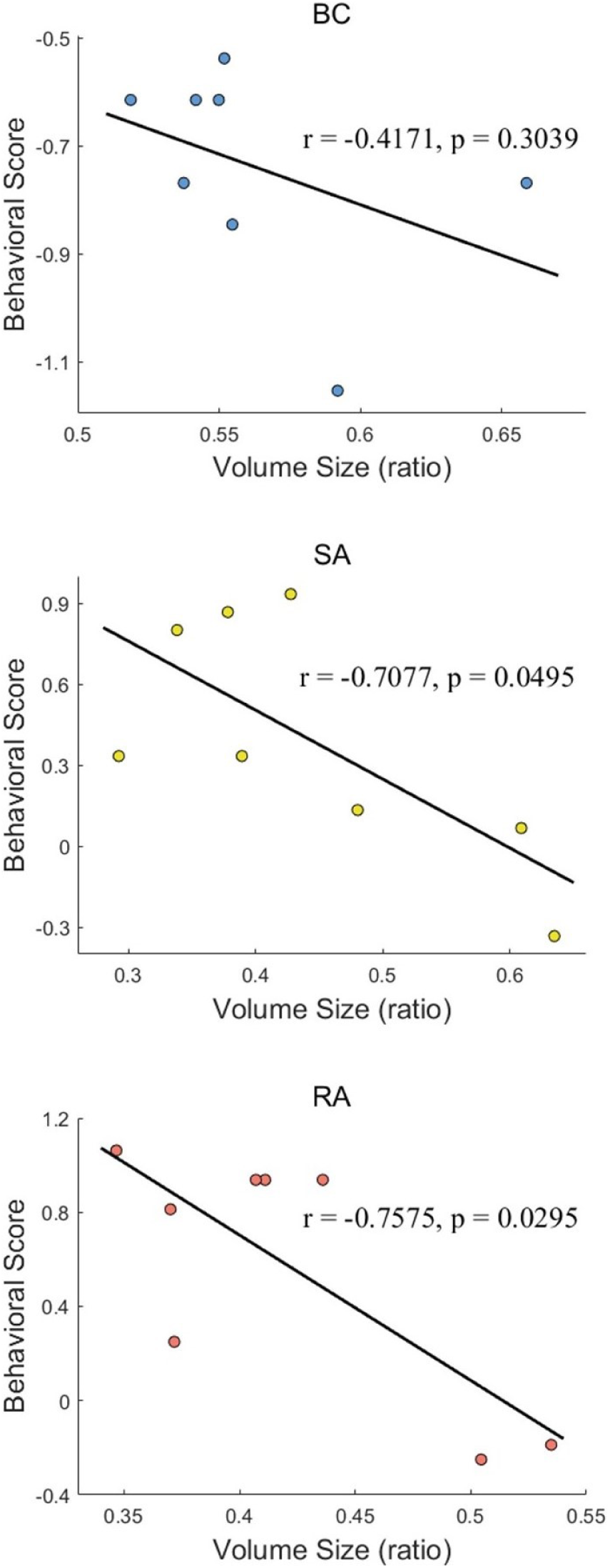
(Extended) Correlation between the hypermetabolic level and behavioral outcomes. Scatter plots of the hypometabolic volume sizes (ratio over the whole volume size) in the ipsilateral right hemisphere (Fig. 1c right) and the mean of the normalized behavioral scores (Fig. 4c right) from day 3 to day 10. Data from the BC, SA, and RA groups are presented from top to bottom.

**Table 1.** (Extended) Seven-point grading scale for the balance beam walking test.

| Scores | Criteria |
| --- | --- |
| 1 | The rat is unable to move or directly falls down from the beam |
| 2 | The rat moves but falls down before the first quarter point of the beam |
| 3 | The rat falls down between the first quarter and halfway point of the beam |
| 4 | The rat falls down from a position between the halfway and three-quarter points of the beam |
| 5 | The rat falls down after the three-quarter point of the beam |
| 6 | The rat crosses the beam but with foot slips |
| 7 | The rat crosses the beam normally |

### Acupuncture modulation of motor control pathways

Walking or locomotion, in both humans and animals, is a complex motor behavior that involves sensory signals about the environment and transmission of motor commands from cortical regions in the brain to neurons located in the spinal cord [37]. In this study, our findings reveal that acupuncture yielded the most significant benefits in the motor cortex and primary somatosensory cortex. Intriguingly, these two regions have been identified as crucial components involved in motor control. Existing studies have provided evidence of the close association between the motor cortex and S1 in the regulation of mammalian locomotion [38], particularly in relation to the somatosensory representations of the forelimb and hindlimb in S1 [39].

In addition to the motor and somatosensory cortex, we have also observed prompt metabolic restoration in regions such as the visual cortex (Extended Fig. 6a), parietal association cortex (Extended Fig. 7a), and the initial segment of the spinal cord (Extended Fig. 8a) in rats from the RA group. In the visual cortex, specifically the primary visual area (V1), the RA group consistently exhibited smaller hypometabolic volumes throughout the treatment period (Extended Fig. 6b). The secondary visual area (V2), which consists of lateral and medial components adjacent to V1, demonstrated less significant efficacy of RA compared to SA and BC (Extended Fig. 6c). However, the early rehabilitation effect was still notable. The recovery area was observed to initiate from the periphery and extend towards the inner cortical layer (Extended Fig. 6d and e). Notably, the results in the visual cortex did not provide evidence supporting the efficacy of SA in reducing hypometabolism compared to the BC condition. The parietal association cortex comprises three sub-regions: the lateral area (LPtA), medial area (MPtA), and posterior area (PPtA), situated at the interface between the somatosensory cortex and visual cortex. Enhanced metabolism and facilitated rehabilitation were observed across all three (Extended Fig. 7b-d). Hypometabolic regions also exhibited inward recovery, progressing from the periphery towards the core (Extended Fig. 7e-g). Only a limited segment of the spinal cord was captured within the field of view (FOV) of the brain imaging. However, within this region, a substantial and early recovery was observed in the RA group (Extended Fig. 8b), with the hypometabolic volume reaching zero during the mid-term of the experimental period (Extended Fig. 8c).

**Fig. 6.**
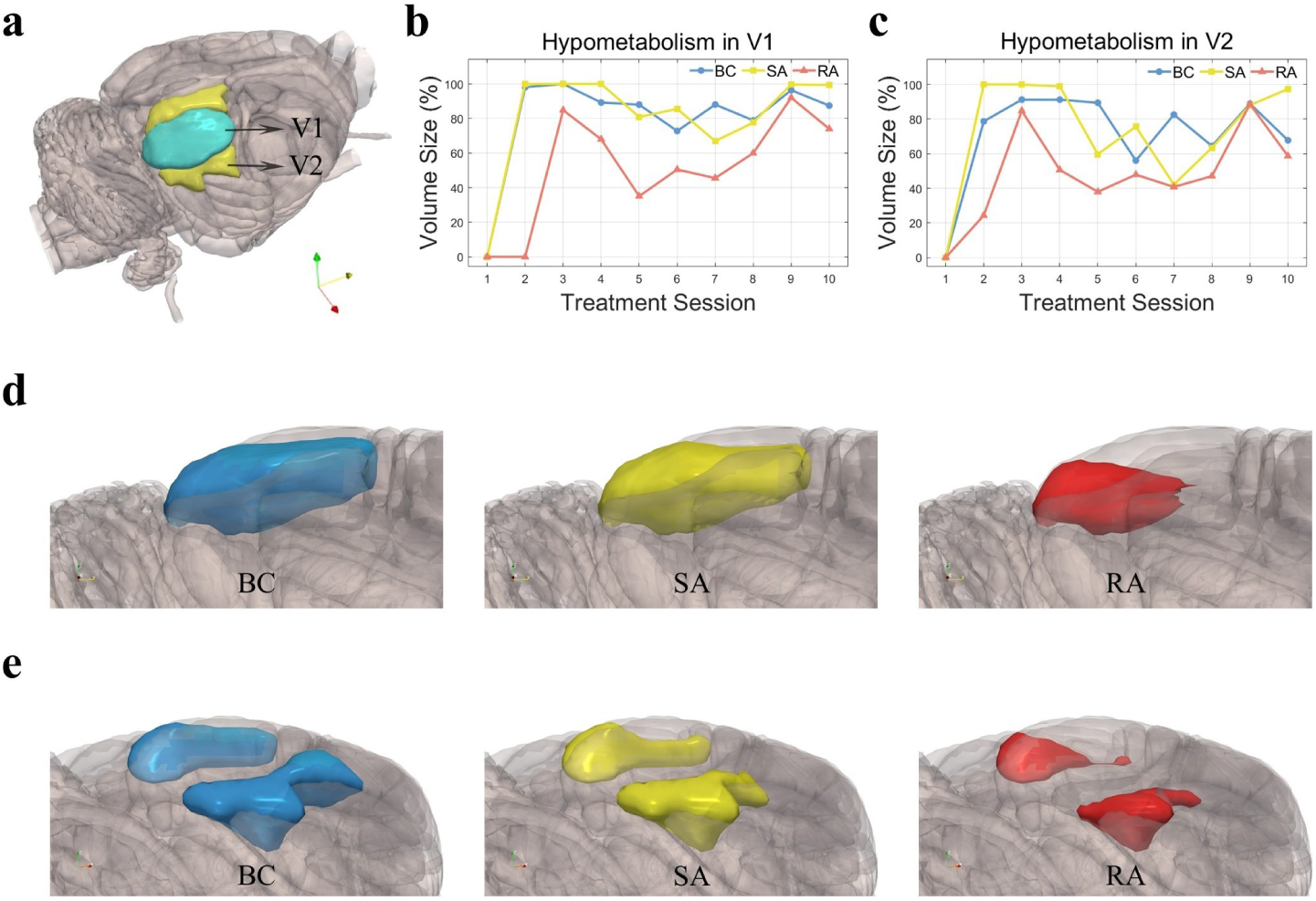
(Extended) Hypometabolic volumes in the visual cortex. **a**, The locations of the primary visual cortex (V1) and secondary visual cortex (V2) in the 3D atlas. **b**, **c**, The ratio of voxels exhibiting hypometabolism in the ipsilateral V1 (**b**) and V2 (**c**) over their original volume sizes, respectively. Calculations and sample sizes are the same as those described in Fig. 2c. **d**, The locations of hypometabolic volumes in the ipsilateral V1 on day 5. Results shown from left to right represent the BC, SA, and RA groups, respectively. **e**, The locations of hypometabolic volumes in the ipsilateral V2 on day 5.

**Fig. 7.**
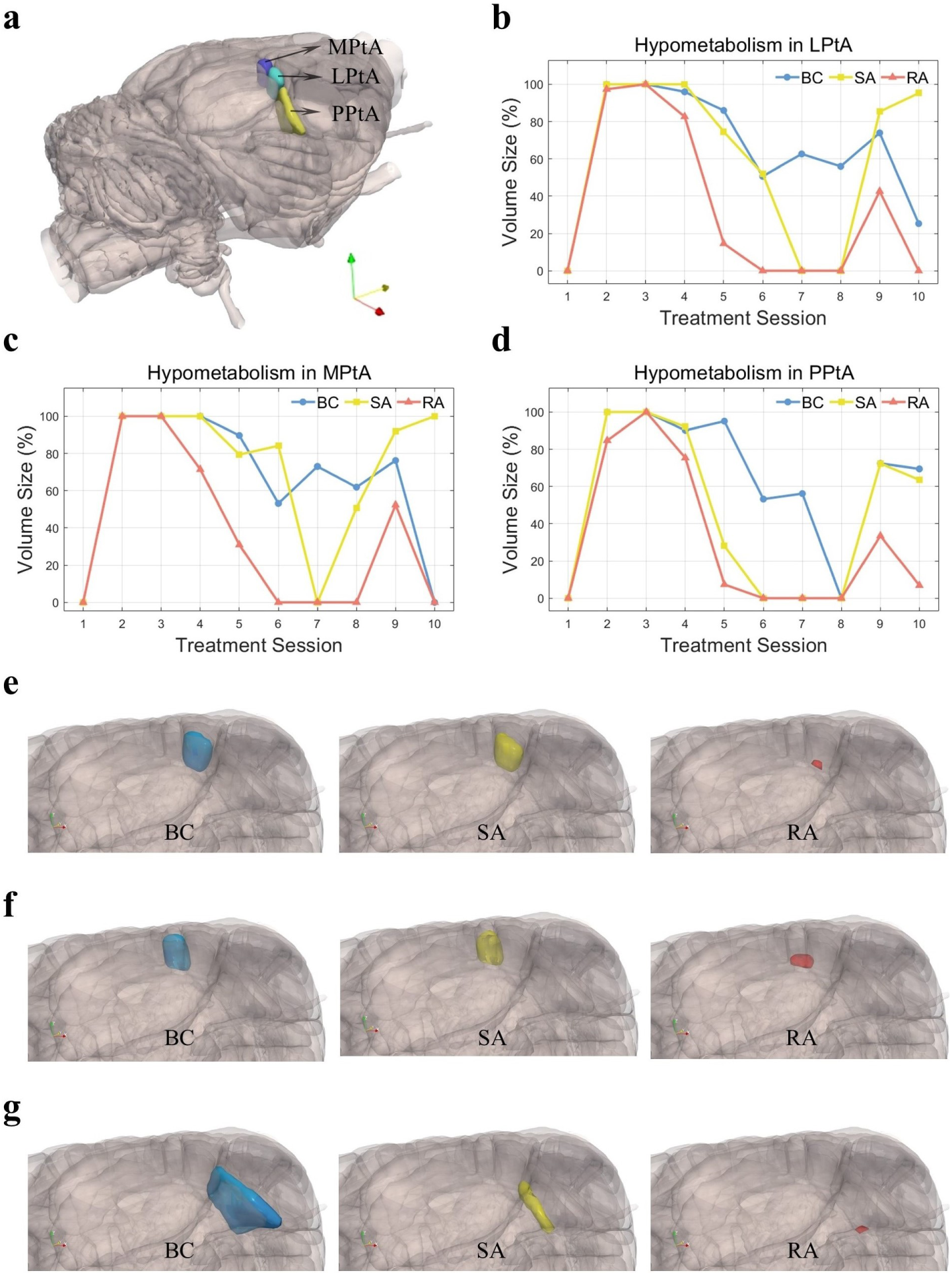
(Extended) Hypometabolic volumes in the parietal association cortex. **a**, The locations of the parietal association cortex: lateral area (LPtA), medial area (MPtA), and posterior area (PPtA) in the 3D atlas. **b** - **d**, The ratio of voxels exhibiting hypometabolism in the ipsilateral LPtA (**b**), MPtA (**c**) and PPtA (**d**) over their original volume sizes, respectively. Calculations and sample sizes are the same as those described in Fig. 2c. **e**, The locations of hypometabolic volumes in the ipsilateral LPtA on day 5. Results shown from left to right represent the BC, SA, and RA groups, respectively. **f**, The locations of hypometabolic volumes in the ipsilateral MPtA on day 5. **g**, The locations of hypometabolic volumes in the ipsilateral PPtA on day 5.

**Fig. 8.**
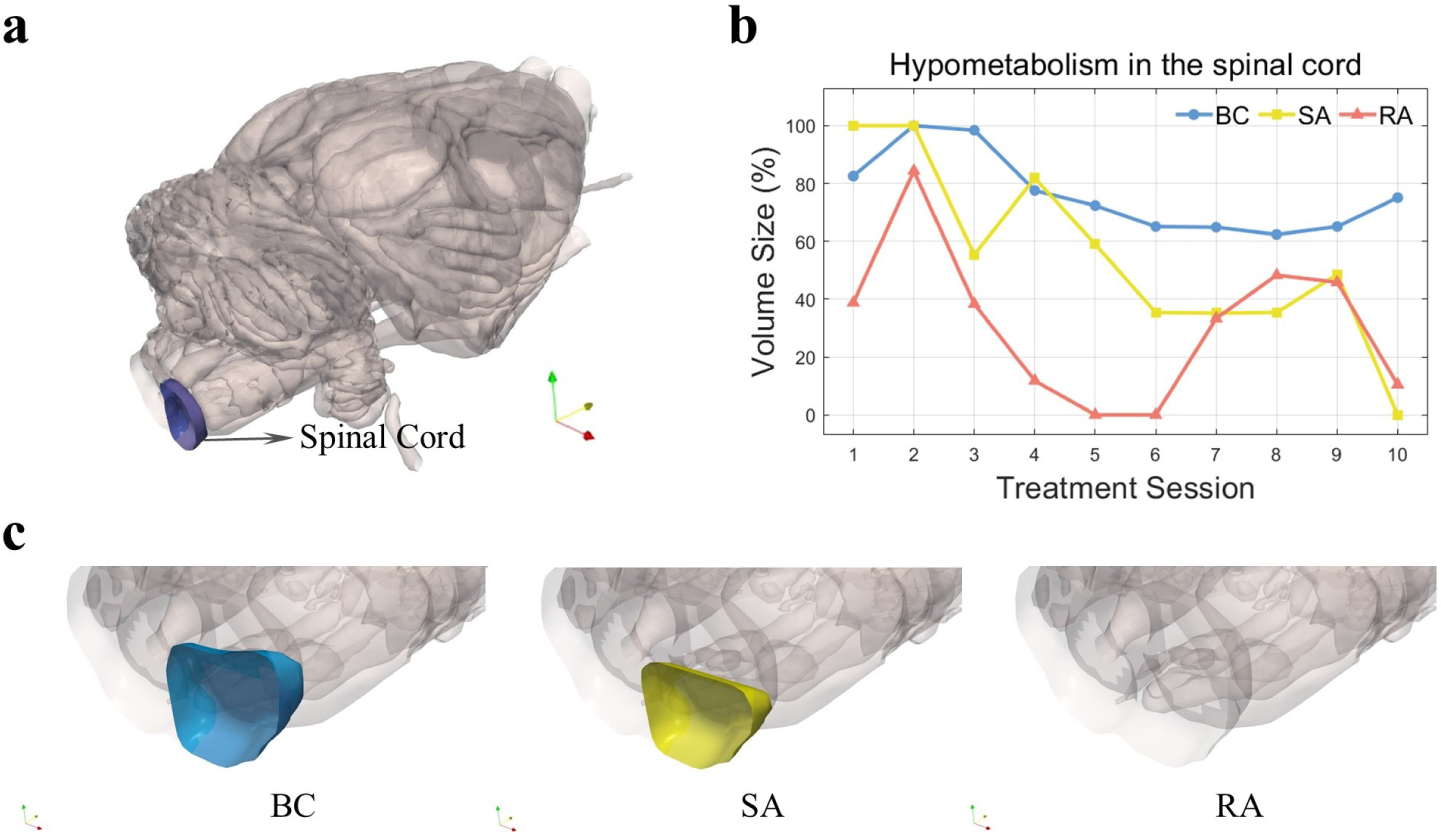
(Extended) Hypometabolic volumes in the spinal cord (partial structure). **a**, The location of a segment of the spinal cord included within the 3D atlas. **b**, The ratio of voxels exhibiting hypometabolism in the corresponding spinal cord over its original volume size. Calculations and sample sizes are the same as those described in Fig. 2c. **c**, The locations of hypometabolic volumes in the ipsilateral spinal cord on day 5. Results shown from left to right represent the BC, SA, and RA groups, respectively.

The primary visual cortex plays a vital role in locomotion by generating predictions of visual flow based on motor commands through a top-down projection from M2 and the anterior cingulate cortex [40]. Further, rats with lesions in the parietal association cortex have shown a significant deficit in behavioral tasks, highlighting the crucial role of this area in spatial orientation and movement control [41]. In our study, the beneficial effects induced by acupuncture in these two regions show less consistency when compared to those in M1, M2, and S1. However, this may be because they are situated at a greater distance from the two acupoints in comparison to the motor and somatosensory areas, potentially indicating a hierarchical pathway of treatment efficacy that propagates from the frontal lobe to the parietal-temporal and occipital lobes (Extended Fig. 9a). Together with the spinal cord, they form vital components that regulate locomotion. The behavioral test scores quantitatively demonstrate the enhancement of motor ability in the rats following acupuncture and the superior recovery effects of acupoint stimulation compared to non-acupoint stimulation. This validation further supports the imaging findings from this study.

**Fig. 9.**
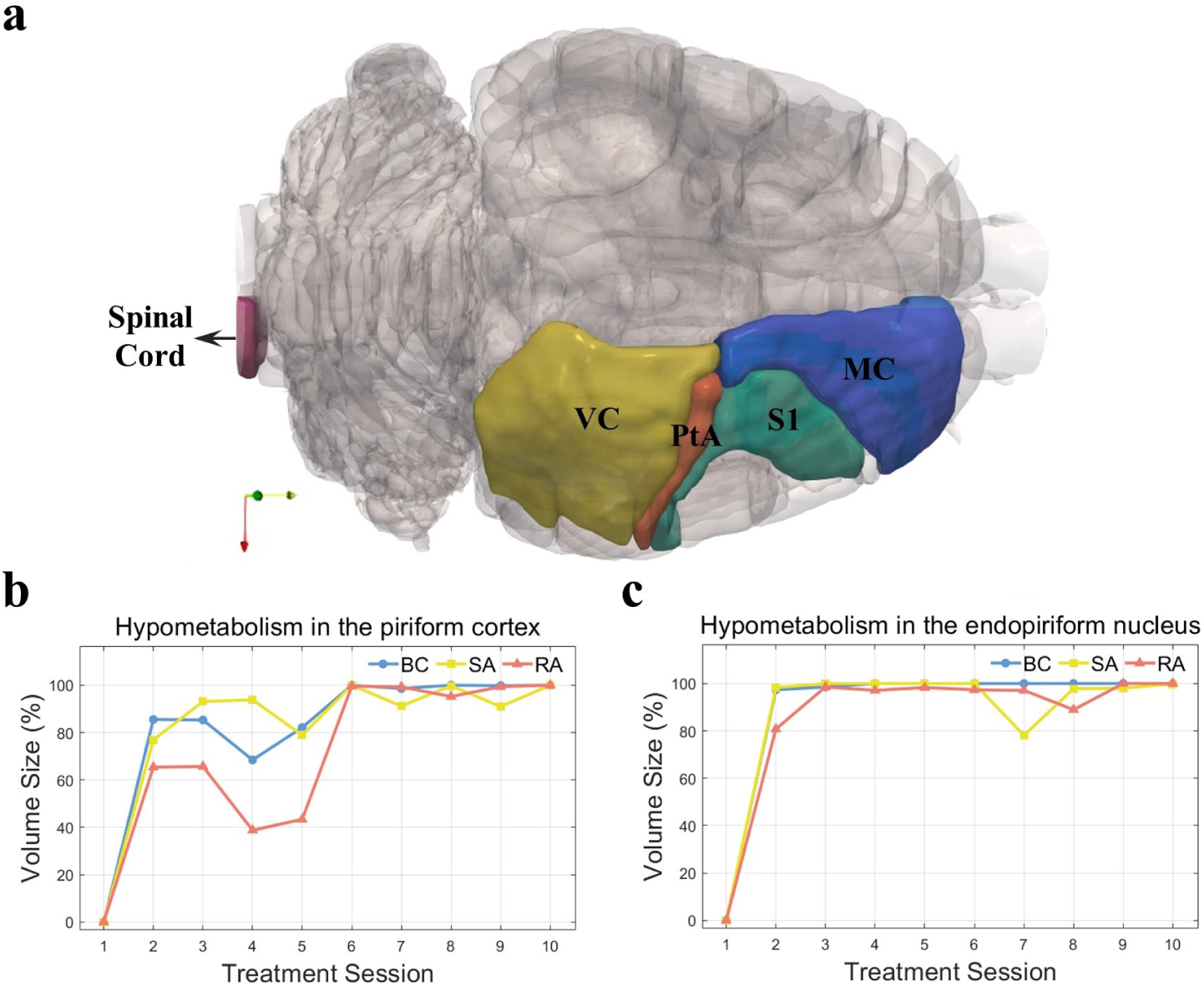
(Extended) Brain regions with and without treatment effect. **a**, Four cortical regions within the motor pathway, together with the spinal cord, that demonstrated promoted recovery from RA. MC: motor cortex; PtA: parietal association cortex; VC: visual cortex. **b**, **c**, The ratio of voxels exhibiting hypometabolism in the piriform cortex (**b**) and endopiriform nucleus (**c**) over their original volume sizes, respectively. Calculations and sample sizes are the same as those described in Fig. 2c. These two regions have lesions evident in the MRI scans (Fig. 4a) and infarction observed through TTC staining (Fig. 4b).

The regions identified as the ischemic core are marked as lesions in the MRI scans (Fig. 4a) and infarctions in the TTC staining results (Fig. 4b). They are thus supposed to be beyond the salvageable reach of acupuncture treatment. Hypometabolic volumes in the piriform cortex (Extended Fig. 9b) showed a facilitated rehabilitation response in the RA group during the initial five treatment sessions, indicating its viability after MCAO. However, starting from day 6, a sudden escalation of hypometabolism was observed in all three groups, and the hypo-uptake volume size remained consistently around 100% until the end of the study. The endopiriform nucleus exhibited clear evidence of being an ischemic core, with the potential development of infarction occurring on the second day following the MCAO procedure (Extended Fig. 9c).

Considering that the therapeutic effect of acupuncture on stroke may be relatively subtle, after conducting hypothesis tests on different voxels, we adopted the BH procedure to address the multiple comparison problem and improve statistical power. To mitigate the risk of obtaining an early recovery pattern as a result of false positives, we conducted additional analyses employing the Bonferroni correction. The sizes of the hypometabolic volumes obtained using this conservative strategy still demonstrated a distinct early recovery associated with RA in M1, M2, S1FL, S1HL, LPtA, MPtA, and the spinal cord (Extended Fig. 10).

**Fig. 10.**
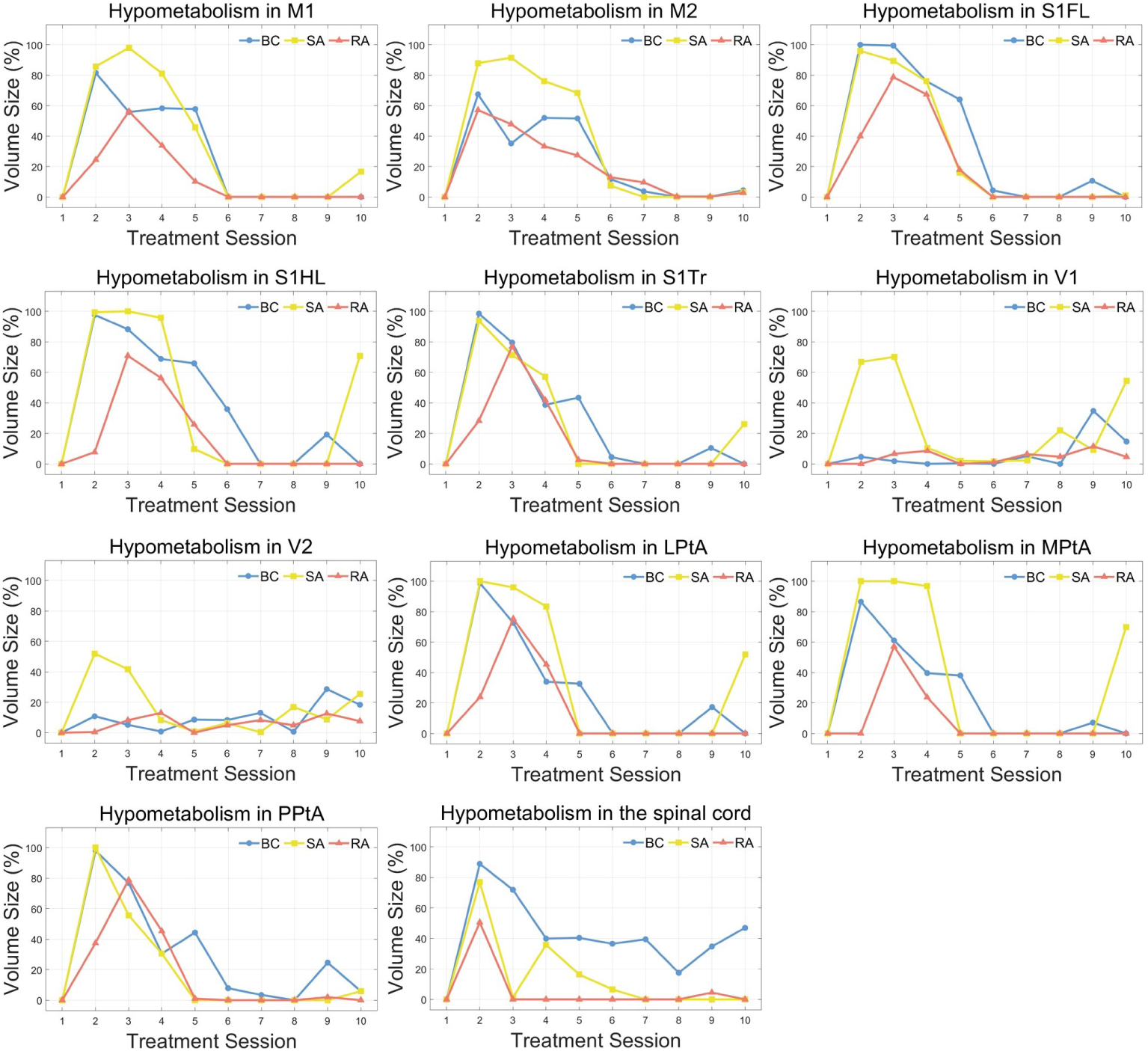
(Extended) Hypometabolic volume sizes using the Bonferroni correction. Voxels demonstrating a statistically significant decrease in FDG uptake were identified following the same calculations as described in Fig. 2c, but with the Bonferroni correction (*p* = 0.01) for family-wise error control instead of the BH procedure.

## Conclusion

This study presents quantitative evidence regarding the efficacy of acupuncture in facilitating recovery from stroke. Utilizing microPET imaging to measure metabolic levels in various brain regions, we conducted a comparative analysis involving rats that received no treatment, rats that underwent acupuncture at GV20 and GV26 acupoints, and rats that underwent needling at sham acupoints. The findings revealed that manual needling at both acupoints and non-acupoints demonstrated a therapeutic effect by reducing the size of the hypometabolic volume. Furthermore, in regions associated with motor control, real acupuncture exhibited an earlier reduction in hypometabolism compared to sham acupuncture. These findings were consistent with the outcomes of a balance beam walking test, which assesses motor deficit. Overall, the results suggest the existence of a neural pathway through which acupuncture modulates the rehabilitation of motor function.

## Methods

### Animals

Female Sprague-Dawley rats (180–280 g) were included in this study. All rats were housed in a temperature-controlled room that followed a 12-h light/dark cycle, with lights on from 8.00 a.m. to 8.00 p.m. and lights off from 8.00 p.m. to 8.00 a.m.. All animals were provided with unrestricted access to food and water during the experiments, with the exception of an overnight fast preceding each scanning session. For all rats, we applied the same experimental procedure, comprising brain surgery, anesthesia, microPET radiotracer preparation, microPET imaging, and functional assessment. The only difference lay in the treatment step. Experiments were approved by the Animal Research Committee of the School of Medicine, Zhejiang University. All the procedures complied with the ARRIVE guidelines and were carried out following the National Institutes of Health Guide for the Care and Use of Laboratory Animals (NIH Publications No. 8023, revised 1978).

### Focal ischemia

Before the cerebral ischemia was induced, rats were placed on a heating pad with their temperature maintained and body weight measured. Then the rats were anesthetized with isoflurane (2% *−* 3%). The ischemic model was performed via middle cerebral artery occlusion (MCAO) [42] with craniotomy and electrocoagulation. First, the skin was cut with scissors, causing the temporalis muscles to retract, revealing zygomatic and squamosal bones underneath. A high-speed drill was employed to generate a tiny aperture (2-3 mm) at the junction of the squamosal and zygomatic bones. The middle cerebral artery (MCA) in the right hemisphere was recognized and exposed through the technique of dura retraction with the aid of ophthalmic scissors and forceps. Subsequently, the MCA was permanently occluded through the application of bipolar electrical coagulation. Sterilized absorbent cotton was utilized to pack the surgical area, and the skin was closed using sutures. Following the MCAO procedure, the rats were randomly allocated to one of the three treatment groups. Rats that died during the experiment were excluded from the data analysis.

### Magnetic resonance imaging

Magnetic resonance imaging (MRI) was conducted 2 h after the induction of stroke to validate the brain injury resulting from the successful ligation of MCA. The data were acquired using a GE Signa Excite 1.5T MRI scanner located in the Sir Run Run Shaw Hospital of Zhejiang University. The scans were converted from DICOM (Digital Imaging and Communications in Medicine) format into Analyze format using MRIcron (https://www.nitrc.org/projects/mricron) and then aligned to the Waxholm Space atlas in the statistical parametric mapping software (SPM12, Wellcome Department of Imaging Neuroscience, London, United Kingdom). The registration to the atlas enabled the identification of the ischemic lesions in an acute stage (Fig. 4a). It also ensured the successful induction of brain ischemia via MCAO surgery. Due to the large number of samples involved in the experiment, only randomly selected rats (*n* = 16) underwent MRI scans.

### Localization of infarct tissues

A histochemical evaluation of brain infarction was performed to confirm the success of MCAO and identify the location of the infarction. 2,3,5-triphenyltetrazolium chloride (TTC) staining was conducted on randomly selected rats (*n* = 9 from the BC group) 24 h after MCAO. The animals were sacrificed, and their heads were immediately removed and sliced into 2 mm thick sections. The coronal brain sections were then incubated in a 2% TTC solution at 30*^◦^C* for 20 min. The presence of white-colored areas indicated infarct lesions (Fig. 4b), thereby confirming the successful induction of ischemia.

### Needling

Rats were anesthetized with isoflurane (2% *−* 3%), and their body temperature was maintained using a heating pad. To rule out the influence of electrical stimulation on stroke recovery, manual acupuncture was administered instead of electroacupuncture. The acupuncture procedure was identical for subjects in the RA and SA groups, with the exception of the needling targets. The procedure involved a pair of stainless filiform needles (0.18 *×* 9 mm, Hwato). In the RA group, the needles were simultaneously inserted into GV20 (5.0-10 mm deep) and GV26 (2.0-3.0 mm deep). Following insertion, the needles were twisted for 1 min at a frequency of 2.0 *−* 2.5 Hz, with an even back-and-forth amplitude of approximately 180 degrees. Then, the acupuncturist stopped the procedure for 4 minutes. This cycle was repeated six times in total, taking a total duration of 30 min. In contrast, for animals in the SA group, needles were inserted into two sham acupoints located 5 mm to the right of GV20 and GV26, respectively (Fig. 1b). The initial real or sham acupuncture was carried out 4 h after the MCAO surgery. The treatment for each animal commenced on a Monday and continued until Friday of the second week, spanning a duration of ten consecutive working days. In order to minimize differences in treatment effects among animals due to variation in needling techniques, all acupuncture treatments were performed by the same experienced acupuncturist.

### FDG microPET imaging

MicroPET imaging was conducted for each rat prior to MCAO, 0.5 h after MCAO, and following each acupuncture session. Owing to the extensive sample size involved in the study, the baseline imaging and MCAO procedure were not performed on the same day. Instead, the baseline images were obtained on the Friday preceding the induction of cerebral ischemia (the ensuing Monday). Rats were anesthetized using isoflurane (2% *−* 3%) prior to the baseline imaging; however, anesthesia was not employed for the MCAO and acupuncture imaging as the animals were already anesthetized before the MCAO and acupuncture procedures. Thirty minutes post-administration of FDG with a dose of 1 mCi through the tail vein, each rat was positioned within the microPET machine and subjected to a 15-min scan. The imaging session was carried out immediately after the MCAO procedure and each treatment trial. However, due to the 30-min resting period required for FDG uptake, the gathered data represented the glucose uptake levels 0.5 h post-cerebral ischemia or acupuncture. Animals in the BC group did not receive any treatment, but they also underwent a microPET scan on each treatment day to reveal the natural rehabilitation from stroke without any intervention.

All microPET data were collected in 3D mode using a microPET R4 scanner (32 detector rings; animal aperture, 120 mm; Field of view (FOV), 78 mm; CTI Concorde Microsystems, LLC) situated at the Medical PET Center, Zhejiang University. The spatial resolution of the microPET camera is 1.9 mm full width at half maximum (FWHM) in the transverse plane and 1.88 mm FWHM in the axial plane. Each animal was placed in the transaxial position with its head in the FOV of the microPET R4 scanner. A 15-min static FDG microPET image was acquired, and the raw data were reconstructed as 128 *×* 128 *×* 63 volumes with a voxel size of 1 *×* 1 *×* 1 mm^3^. The reconstruction algorithm is based on the maximum a posteriori method with 18 iterations and a resolution parameter (*β*) of 0.5, as provided by the supplier. The reconstructed data were saved in Analyze format and preprocessed in SPM12, following the pipeline described in the previous work [43]. In summary, the images were first resized based on an in-house FDG-microPET template specifically designed for Sprague-Dawley rats. Second, each image was approximately registered to the template using the realign function in SPM. Third, the intracerebral volume was segmented and registered to the template using the SPM co-register function. Fourth, warping parameters were estimated to map the image onto the template using the normalise function in SPM. Considering the gross pathology observed in the ipsilateral right brain, only the left hemisphere was utilized for the estimation of the normalization parameters. Finally, the images were smoothed using the SPM smooth function with a Gaussian kernel of 2 *×* 2 *×* 4 mm^3^ FWHM. The resulting image was stored in an 80 *×* 108 *×* 63 volume with a voxel size of 2 *×* 2 *×* 2 mm^3^. Furthermore, a volumetric atlas of the rat brain, known as the Waxholm Space atlas (https://www.nitrc.org/projects/whs-sd-atlas), was spatially normalized to conform to the same coordinate space [31]. This atlas comprises 222 brain structures, including a comprehensive parcellation of the cerebral cortex.

### Image analysis

To reduce the variability of voxel intensities across FDG microPET images, intensity normalization was performed using the contralateral left brain. Voxel intensities from each image were divided by the mean intensity over its left hemisphere, and the resulting intensities represented relative metabolic values. These values had smaller variability compared to the absolute cerebral metabolic rates of glucose (CMRglc) and did not require blood sampling [44]. Following intensity normalization, difference images were generated by subtracting the MCAO scan from each treatment scan. The rationale for using the MCAO scans instead of the baseline scans was that we believed the metabolic changes resulting from focal ischemia would be more substantial than the metabolic recovery induced by acupuncture. Consequently, the difference between the baseline and treatment scans would be primarily influenced by the MCAO effect, and the treatment differences among the three groups would be concealed.

Paired *t*-tests were utilized to identify hypo- and hyper-uptake voxels with one-tailed tests. Here the decreasing side denotes hypometabolism and the increasing side denotes hypermetabolism. P-values were computed for each voxel and sorted in ascending order. The reordered p-values were then compared one by one with a critical value using the Benjamini-Hochberg (BH) procedure [30]. A threshold was established to define significant voxels based on a controlled FDR level of 0.01. The hypometabolic/hypermetabolic volume size was determined by calculating the number of significant voxels within the original volumes.

### Assessment of motor deficit

A balance beam walking test was performed to evaluate the neurological deficit of rats from each group (Fig. 1a). A wooden beam (100 cm in length, 3.7 cm in diameter, and 20 cm off the floor) was placed horizontally for the assessment. In view of temporal limitations, the behavioral test was conducted only on rats chosen randomly from each group, starting from the first day of treatment to the last day. To ensure the objectivity of the assessment, the investigator was unaware of the ID and treatment group affiliation of each rat being tested. The motor deficit was evaluated based on a seven-point grading scale (Extended Table 1) with reference to a previous study [45].

### Statistical analysis

Results are expressed as mean *±* standard error of the mean (SEM). Statistical analyses were accomplished using MATLAB R2022b software. After MCAO, animals were randomly assigned to treatment groups via a three-number lottery draw. When computing significant voxels from image data, a one-tailed paired *t*-test was employed, with FDR controlled at a level of *q* = 0.01 using the BH procedure. All the data under *t*-tests or ANOVA were checked for normality with the Kolmogorov-Smirnov test. P-values less than 0.05 were considered statistically significant.

## Data availability

All data included in this paper are available at: http://yulab.ust.hk/acupuncture/.

## Code availability

The MATLAB code used in the analysis of the current study is available at: http://yulab.ust.hk/acupuncture/.

## Acknowledgements

This work was supported in part by the internal fund 3030-009 and BGF.001.2023 from HKUST, T12-101/23-N (RGC), R4012-18 (RGC), 7015-23G (RGC), and MHP/033/20 (ITC) from the Hong Kong S.A.R. Government of China, the National Key Technology Research and Development Program of China (No: 2017YFE0104000), and the National Natural Science Foundation of China (No: 61525106, U1809204). We thank Tania Wilmshurst for proofreading the manuscript. We thank the Generic Diagramming Platform (https://biogdp.com/) for providing a tool conducive to scientific illustrations.

## Author contributions

W.Y. and H.L. conceived the study and designed the experiments. H.T., J.L. and X.S. performed the experiments and acquired imaging data. W.H. and C.L. analyzed data. W.H. and W.Y. wrote the manuscript. All authors discussed the results and commented on the manuscript.

## Materials and Correspondence

Correspondence and requests for materials should be addressed to Weichuan Yu.

